# A century-old museum sample reveals a bandavirus with modern day presence in northern European bats

**DOI:** 10.1101/2025.02.28.640763

**Authors:** Udo Gieraths, Jörn Beheim-Schwarzbach, Matthew J. Pickin, Annika Beyer, Lineke Begeman, Bernd Hoffmann, Kore Schlottau, Martin Beer, Rainer G. Ulrich, Thomas Müller, Conrad M. Freuling, Tiina Mauno, Marco van de Bildt, Vera C. Mols, Victor M. Corman, Friedemann Weber, Terry C. Jones, Christian Drosten

## Abstract

Ancient genome sequences provide invaluable insights into the origin and evolution of viral pathogens, offering a broader temporal perspective that extends well beyond the limited timespan of clinical data, which typically covers at most a few decades. Whereas ancient viral DNA is relatively frequently recovered, ancient viral RNA genomes are scarce due to the fragile nature of RNA molecules. In this study, we explored the feasibility of detecting ancient viral RNA in ethanol-preserved bat samples from a museum collection. We not only detected viral genome fragments but also recovered the coding-complete genome of a bandavirus (species *Bandavirus zwieselense*, family *Phenuiviridae*, order *Hareavirales*, class *Bunyaviricetes*) from a Common pipistrelle (*Pipistrellus pipistrellus)* bat collected in northern Germany in 1919.

To investigate the modern distribution of *Bandavirus zwieselense*, we screened bat organs collected in Germany and the Netherlands via RT-qPCR, identifying modern counterparts in nine Common Pipistrelle (P. pipistrellus) bats (collected between 2010 and 2018), and one Serotine bat (*Eptesicus serotinus*, collected in 1999). The resulting genomic data enabled us to map phylogenetic relationships within this previously uncharacterized virus species (*Bandavirus zwieselense*) and estimate the timeframe for the most recent common ancestor. Additionally, we performed functional analysis of the S segment encoded nonstructural (NSs) protein of *Bandavirus zwieselense* in human HEK-293T cells, demonstrating its ability to block interferon induction, a characteristic also observed in the related human-pathogenic *Severe fever with thrombocytopenia syndrome virus (*SFTS*V, species Bandavirus dabieense)*.

This study demonstrates the feasibility of recovering and characterizing viral genomes from ethanol-preserved ancient RNA material, underscoring the significant potential of museum collections to contribute to the understanding of RNA virus evolution.

## Introduction

In recent years, significant progress in the study of virus evolution has been driven by ancient exogenous viral sequences discovered in the context of archeological studies^1–7^. Findings of human DNA viruses, such as *Hepatitis B virus*^*1*,*4–7*^, *Parvovirus B19*^*2*^ and *Variola virus*^*3*^, with sequences dating back thousands of years, have provided unprecedented insights into the evolutionary dynamics of DNA viruses.

In contrast, research on ancient RNA viruses has primarily focused on viral genomes recovered from the past century in medical collection or outbreak-associated samples, such as *Human immunodeficiency Virus 1* from 1959^8^ and 1960^9^, *Measles virus* from 1912^10^, and *Influenza A virus* from the 1918 pandemic^11^. Exceptions include the *Barley Stripe Mosaic Virus* genome recovered from ~750-year-old grain^12^. Although not of viral origin, small RNA fragments were detected in 3,400-year-old seeds^13^ and the oldest recovered RNA was from permafrost-preserved liver tissue of a canid dating back 14,300 years^14^. These exceptional cases are largely confined to environments that naturally limit RNA degradation, such as permafrost or the protective conditions within plant seeds.

RNA degradation is also mitigated by human-made preservation in the form of formalin-fixed human tissues. Pathology collections hosting these tissues have been successfully used to recover ancient viral RNA^10,15^. Complementing these human samples, natural history museums house extensive collections of well-preserved animal specimens. Sampled from diverse geographic regions, these specimens offer a valuable resource for tracking zoonotic transmission—including cases where the ancient host species is now extinct. Wet collections, which constitute a significant portion of museum specimens^16^, may be particularly promising for RNA virus preservation, as specimens are typically stored in 70% ethanol, an environment that limits RNA degradation via inactivation of RNases^17^. Ancient viral genomes provide a unique opportunity to reconstruct transmission routes^18^, gain deeper insights into viral evolution^10,15^ and can even enable full virus reconstruction through reverse genetics^19^. Screening ancient specimens can reveal previously unknown viruses that lack modern counterparts. In such instances, ancient genomes can serve as a reference to identify related contemporary viruses, allowing researchers to study viral evolution across extended timescales.

Bandaviruses (genus Bandavirus, family *Phenuiviridae*, order *Hareavirales*, class *Bunyaviricetes)*, are characterized by a segmented RNA genome consisting of three segments: the large (L) segment encoding the RNA-dependent RNA polymerase (RdRp), the medium (M) segment encoding the glycoprotein precursor Gn/Gc, which is post-processed into Gn and Gc, and the small (S) segment encoding the nucleocapsid (N) and nonstructural (NSs) proteins. Multiple studies have demonstrated immunomodulatory functions of the bandavirus NSs protein^20–24^. Among bandaviruses, *severe fever with thrombocytopenia syndrome virus (*SFTSV), also named *Dabie bandavirus (*species *Bandavirus dabieense)*, was identified in 2009 as the causative agent of a tick-borne disease in China^25^. Between 2011 and 2021, 18,902 cases were reported in China, with a case fatality rate of 5.11%^26^.

SFTSV is known to strongly suppress interferon (*IFN*) induction in host cells by sequestering key immune regulatory factors (e.g. Interferon regulatory factor 3, IRF3 and TANK-binding kinase 1, TBK1) in viral inclusion bodies via its *NSs* protein^20–22,27^. Even though SFTSV is suspected to be primarily transmitted via ticks ^28,29^, rare instances of human-to-human transmission are known^30–33^. Since its initial identification, SFTSV has been reported across Asia^34–39^, and related viruses such as *Heartland virus* (*HRTV*)^40^, *Guertu virus*^*41*^ and *Kinna virus*^*42*^ have been discovered. Collectively, these SFTSV-like bandaviruses form a distinct clade within their genus, and RT-PCR-based or serological evidence indicates their presence in a range of domestic and wild mammals^43–50^, suggesting a broad host tropism. However, two additional viruses within the SFTSV-like clade have been identified exclusively in bats: *Malsoor virus*, isolated from two fruit bats, Leschenault’s rousette (*Rousettus leschenaultii*)^51^ in India, and *Zwiesel virus*, detected in five Northern bats (*Eptesicus nilssonii*) in southern Germany^52^.

Here we describe the detection and subsequent investigation of a bandavirus genome found in an ethanol-preserved bat specimen from the wet collection of the Natural History Museum in Berlin.

## Materials and Methods

### Sample Collection and Extraction

#### Natural History Museum

We received partial organ samples (stomach, colon and liver) of 70 bat specimens from the Natural History Museum in Berlin (Supp. Table 3). The specimens were preserved in 70% ethanol and, according to the curators, had not been treated with formaldehyde. Each organ sample was rinsed with Dulbecco’s phosphate-buffered saline (DPBS), cut into cubes of 1-2 mm^3^, and homogenized in DPBS using a Qiagen TissueLyser. The homogenate underwent overnight proteinase K (10 μl) digestion (10-12 h) at 56° C, followed by protease inactivation at 70°C for 10 minutes. After centrifugation the supernatant was transferred to a 2 ml tube and 1 mL of TRIzol™ (Invitrogen) was added, followed by standard RNA extraction according to the manufacturer’s instructions.

#### Recent Bat Samples, Germany

RNA extracts from pooled organs of 1000 bat specimens were obtained as described in Dafalla et al.^53^ and RT-qPCR tested. For specimens that tested positive, individual organ samples (lung, liver, colon) were homogenized in 500 μl of phosphate-buffered saline (PBS, see also Dafalla et al.^53^). A total of 750 μl of Qiagen AL Buffer was then added to 250 μl of homogenate, followed by heat treatment at 75°C for 30 minutes to inactivate the virus. Subsequently, 200 μl of the inactivated homogenate were added to 700 μl Qiagen RLT buffer, and total RNA was extracted using the Qiagen RNeasy kit following the manufacturer’s protocol.

#### Recent Bat Samples, Netherlands

Tissues from lung and liver of 86 Common pipistrelle specimens were obtained as described in Mols et al.^54^ and homogenized for 20 sec in 500 μl viral transport medium (VTM) using 1/4” ceramic spheres (MP Biomedicals), followed by centrifugation at 13,000 rpm for 5 minutes. Subsequently 60 μl of supernatant were added to 90 μl lysis buffer (MagNA Pure 96 External Lysis Buffer, Roche) and 3 μl Equine Arteritis virus was added as internal control. Next RNA was isolated using an in-house developed high-throughput method^55^. If a specimen tested positive, all available other samples of this bat were additionally tested for viral presence and viral load estimates.

### RT-qPCR primers and probe design

Following primers and fluorescein amidite (FAM)-labeled probe targeting the S segment were used: fwd: 5’-GGCTAGGTATTTAAGGATTGCTGC-3’, rev:

5’-GAAGATTGGTCGAAAGTAGCAGTG-3’, probe:

5’-CCATGGCTTGCAGAGATTGCAGGTC-3’. Cycler conditions for RT-qPCR were as follows: 55°C for 20 min, 95 °C for 3 min, 45 amplification cycles at 95 °C for 15 sec, 58 °C for 30 sec and a final cooldown step at 40°C for 30 sec. SuperScript™ III RT/Platinum™ Taq Mix (Invitrogen) was used at 25 μL reaction volume according to the manufacturer’s instructions.

The extracts from the Netherlands were tested using the same primers and probe but Fast virus 1 Step Master mix reagent was used instead: 4xTaqman Fast virus mix 5 μl; 1 μl (10 μM) fwd primer, 1 μl (10 μM) rev primer, 0.5 μl (10 μM) probe, 7.5 μl water, and 5 μl RNA (total 20 μl). Cycler conditions were as follows: 55°C for 5 min, 95 °C for 20 sec, 45 amplification cycles at 95 °C for 3 sec and at 60 °C for 31 sec.

### Library preparation, capturing and sequencing

#### Ancient samples

RNA concentrations were measured using the Qubit™ RNA High Sensitivity Assay Kit (Invitrogen). Because most samples did not have detectable levels of RNA, 10 μL of undiluted extract were used for library preparation with the KAPA RNA HyperPrep Kit (Roche), following the manufacturer’s instructions. To prevent further RNA degradation, the initial fragmentation step was omitted. Libraries were sequenced to various depths on an Illumina NextSeq system.

#### Contemporary samples

A total of 100 ng of RNA was used for library preparation with the KAPA RNA HyperPrep Kit (Roche). Approximately 500 ng of each amplified library underwent viral read enrichment using the xGen™ hybridization capture kit (IDT), with capture probes consisting of interleaving 120 base pair (bp) fragments of the *Zwiesel* (MN823639 - MN823641), *Malsoor* (KF186497 - KF186499), and our ancient *Penzlin* virus genomes. Enriched libraries were then sequenced at varying depths on an Illumina NextSeq system.

### Next-generation sequencing data processing

Sequencing data from multiple runs and organs per specimen were combined, and fastp^56^ (v0.23.2) was used to remove duplicates, filter low-quality reads, and merge overlapping paired-end reads (--merge, --dedup --length_required 30). For virus discovery, the processed reads were aligned against a custom database of viral RNA genomes using bowtie2^57^ (v2.4.2) and the *--very-sensitive* flag. To assemble the ancient (Penzlin) virus genome, reads mapping to host ribosomal RNA were discarded (bowtie2^57^ v2.4.2, default parameters), and *de novo* assembly was carried out in SPAdes (v3.15.5, default parameters)^58^. For recent samples, fastp-processed reads were mapped to our assembled ancient Penzlin reference genome (bowtie2^57^ v2.4.2, *--very-sensitive*), and consensus sequences were generated using SAMtools (v1.21, mpileup -aa -A -d 0 -Q 0)^59^ and ivar (v1.4.3, default parameters)^60^.

### Phylogenetic Analysis

Multiple sequence alignments (MSA) were generated using MUSCLE 5.1^61^ with default settings for the following datasets: (1) coding sequences (CDS) of *Bandavirus zwieselense* (*B. zwiesel)* genomes excluding the potential mosaic strain (assembled from five different viruses) *Zwiesel virus* (MN823639.1), (2) L segment sequences of *Bandavirus* reference genomes (see accession numbers in Fig. 1) together with *B. zwiesel* sequences (Fig. 1), and (3) L segment sequences of all *B. zwiesel* genomes excluding the potential mosaic strain Zwiesel virus (MN823639.1). **Maximum Likelihood phylogeny:** Maximum likelihood (ML) phylogenetic trees were constructed using IQ-TREE multicore version 2.3.6^62^ for MacOS Intel 64-bit. A thorough model search was performed before tree computation, and bootstrap support values were estimated using UFBoot^63^. For the ML tree of all bandavirus sequences (Fig. 1, Supplementary Fig. 3), additional bootstrap support values were calculated using SH-aLRT^64^. The resulting tree images were visualized and exported using FigTree v1.4.4^65^ before final modifications were made in Inkscape v1.4^66^. **BEAST analysis:** Each of the following steps were performed on the L segment of the **ancient+** (including the ancient sample) and **ancient-** (excluding the ancient sample) dataset. TempEst v1.5.3^67^ was used to confirm a temporal signal, followed by inference of dated coalescent trees using BEAST v1.10.4^68^. We selected the HKY substitution model and estimated base frequencies. A strict clock rate was chosen using an exponential prior with mean 0.01 substitutions per site per year. Further, a constant size coalescent tree prior was used. For the root height of the tree, a broad normal prior was chosen with mean 1000 years and standard deviation 1000 years. The Markov chain Monte Carlo analysis was run for 10 million iterations with samples taken every 1000 steps, after discarding the initial 1 million burn-in states. Tracer v1.7.2^69^ was used to confirm convergence and mixing, as well as an effective sample size > 200 for all relevant parameters. Kernel density estimates (KDE) were computed for the *age(root)* and the *clock rate* statistic using Tracer v1.7.2^69^. The KDE plot was exported and modifications of the font and additional legends were done using Inkscape v1.4^66^.

**Fig. 1:**
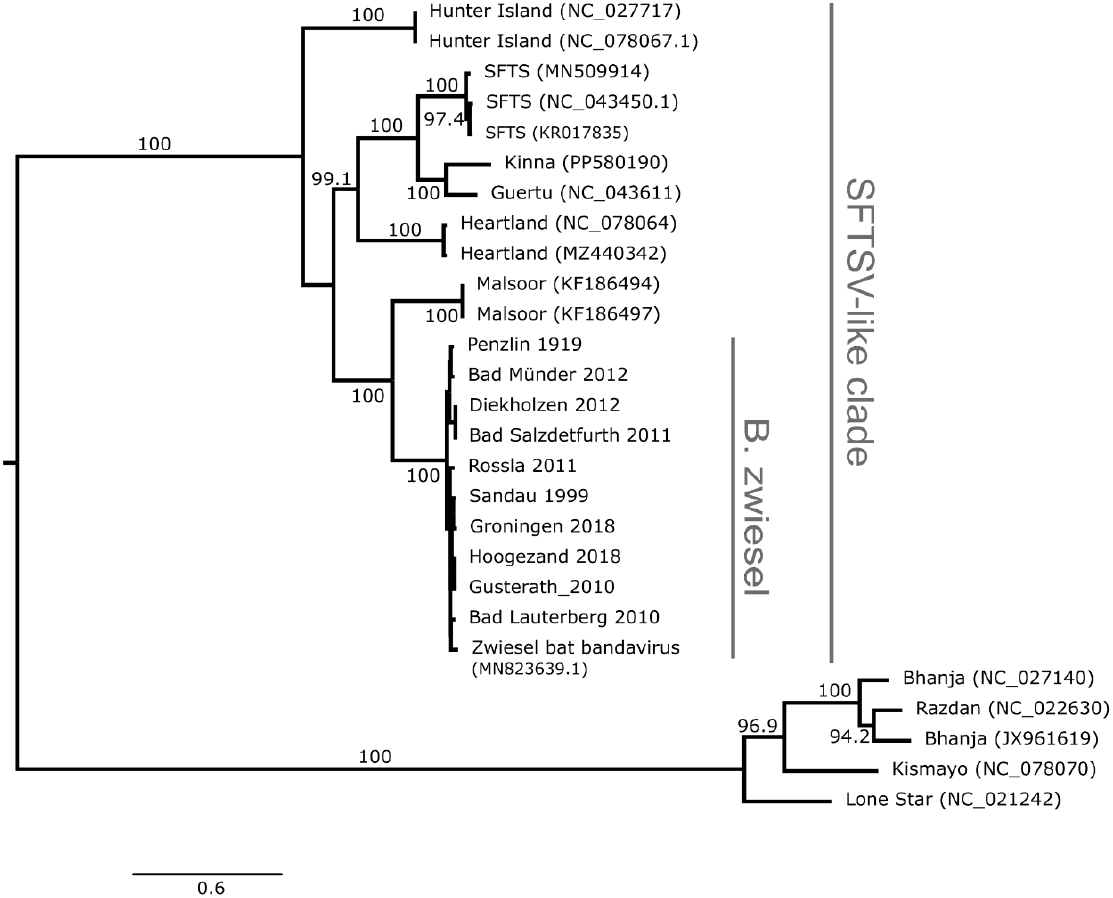
Phylogenetic relationships of strains of the Bandavirus genus on the L Segment. The maximum likelihood tree was computed using IQ-TREE 2.3.6^62^. Rift Valley Fever Virus (NC_014397.1) was used as an outgroup for rooting and subsequently removed for plotting. Bootstrap support values were computed using UFBoot^63^ and only displayed if the support is >= 95%. Within the *B. zwiesel* clade no bootstrap support values are plotted. Supplementary Figure 3 shows the raw tree output, including the outgroup and UFBoot^63^ and SH-aLRT^64^ bootstrap values for all branches.

### Writing of the Manuscript

To enhance clarity, excerpts of this manuscript were rephrased usingChatGPT o1 and ChatGPT-4o.

### Plasmids

Expression constructs for the NSs proteins of SFTSV strain HB29 and *B. zwiesel* strains Penzlin and Bad Lauterberg were synthesized with N- or C-terminal 3×FLAG tag-coding sequences and subcloned into expression vector pI.18 (kindly provided by Jim Robertson, National Institute for Biological Standards and Control, Hertfordshire, United Kingdom). For the two B. zwiesel strains, additional constructs without tags were created. The 3×FLAG tagged HaloTag expression construct was subcloned from a plasmid purchased from Promega into pI.18 with an N-terminal 3×FLAG tag-coding sequence. The luciferase reporter construct *p125-Luc*, expressing firefly luciferase under the control of murine *Ifnb1* promoter was kindly provided by Dr. Takashi Fujita^70^, while the reporter *pRL-SV40*, constitutively expressing Renilla luciferase was purchased from Promega.

### Interferon induction assay

HEK293T cells seeded at a density of 5×10^4^ cells per well in 24-well plates were transfected with 150 ng of *p125-luc*, which expresses firefly luciferase under the control of a *Ifnb1* promoter, alongside 50 ng of *pRL-SV40*, encoding Renilla luciferase under the control of a constitutively active promoter. In a subset of transfection mixes, pI18.MAVS-Strep, which expresses Strep-tagged Mitochondrial antiviral-signaling protein (MAVS), was included to activate the *Ifnb1* promoter. Plasmids expressing the N-, C-terminally 3×FLAG tagged or untagged NSs proteins of *B. zwiesel* or N-terminally 3×FLAG tagged SFTSV were included in varying amounts (30, 100, 300ng). In parallel, an equivalent amount of N-terminally 3×FLAG tagged HaloTag-expressing plasmid was included as a negative control. All transfections were carried out using TransIT-LT1 (Mirus Bio) with total amounts of plasmid DNA being normalised with an equivalent amount of empty pI.18 vector. After 24 hours incubation the cells were processed using a dual-luciferase reporter assay system (Promega) with data read on a Berthold TriStar 2 LB942 Multimode Reader. Firefly luciferase activity was normalized to Renilla luciferase activity, and the resulting values for each construct were then expressed as a percentage of the normalized *Ifnb1* promoter activation observed with the empty vector.

### Immunofluorescence microscopy

Huh7 cells seeded on glass cover slips were transfected with plasmids expressing either the N- or C-terminally 3×FLAG tagged recent *B. zwiesel* NSs, ancient *B. zwiesel* NSs, SFTSV NSs, N-terminally tagged HaloTag as a transfection control or left untransfected.

Transfections were carried out using GeneJammer (Agilent) according to the manufacturer’s specification. After 24 hours, following removal of the growth media, the cells were fixed for 30 mins with 4% paraformaldehyde. Following permeabilization with 0.5% Triton X100 the cells were blocked with bovine serum albumin (BSA) and stained with the mouse anti-FLAG-M2 antibody (Sigma-Aldrich) followed by goat anti-mouse IgG Alexa Fluor 488 (Invitrogen). The coverslips were removed from the growth plate and mounted using 4′,6-diamidino-2-phenylindole (DAPI)-containing mounting fluid (Sigma-Aldrich). Visualisation and documentation were carried out using a EVOS M5000 Imaging System (ThermoFisher).

## Results

### Bandavirus Presence in European Bats

We extracted and Illumina-sequenced RNA from stomach, liver and colon samples of 33 ethanol-preserved bats from various continents and species (Supp. Table 3) provided by the Natural History Museum (NHM) in Berlin. For one of the specimens, a Common pipistrelle bat from Penzlin in Germany, collected in 1919, partial matches of reads against *Malsoor* and *Zwiesel* viruses were found. Additional extractions from liver, lung, stomach and colon samples of this specimen and deeper sequencing allowed us to reconstruct the coding-complete genome via de novo assembly.

Further museum samples (Supp. Table 3) of Common pipistrelle (28 specimens) and Northern bat (9 specimens) from Germany did not reveal any additional findings. To investigate the possible virus presence in contemporary bats and allow for comparison of recent viral genomes with our single ancient finding, we developed an RT-qPCR assay based on the coding-complete ancient genome. Utilizing our RT-qPCR we screened pooled organs from 1000 samples of 20 bat species (see Supp. Table 1) in Germany and 86 liver and lung samples from Common pipistrelle bats in the Netherlands, leading to ten positive findings. Individual organs per positive specimen were tested according to availability (Table 1, Supplementary Table 5). Positive specimens are geographically widespread across northern Germany and the Netherlands. However, sample availability limited the tested locations, hence these findings do not necessarily provide a definitive map of virus distribution in these sampled regions.

**Table 1:**
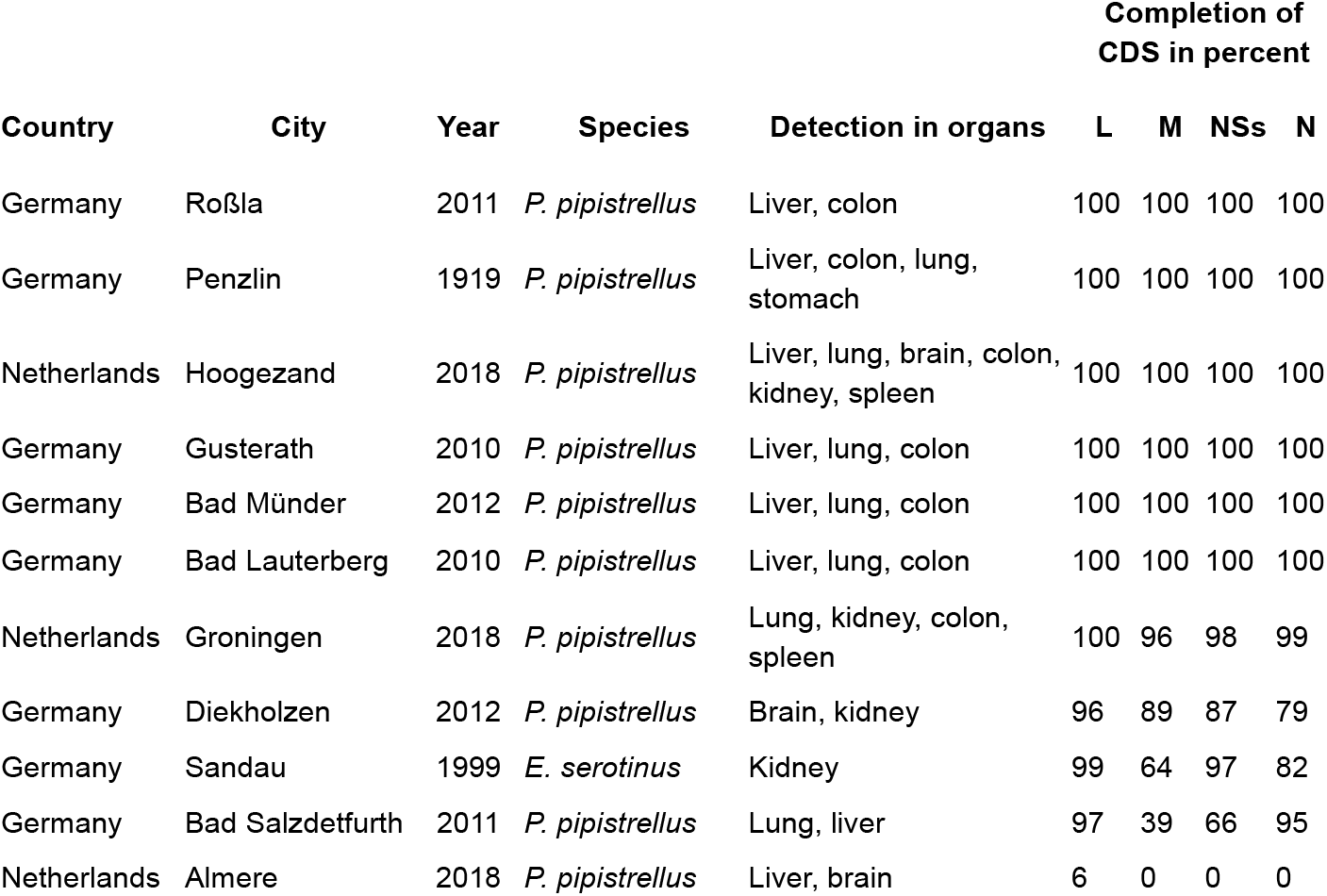
RT-qPCR positive bat specimens. The final column, *completion of CDS in percent*, shows the percentage of the genome of each coding sequence (CDS) that is covered by at least one read, when taking the ancient Penzlin genome as a reference. More sequencing details regarding breadth and coverage can be found in Supp. Table 4.

Sequencing libraries were prepared for the ten positive contemporary samples and enriched using a hybridization capture approach. Taking the ancient Penzlin genome as reference, for nine samples, 96-100% of the coding sequence (CDS) on the L-segment were covered with at least one read (breadth). The coverage varied between datasets and segments. On the L segment we saw an average coverage depth between 131 and 7721 reads across all ten datasets. Details for the other segments with less breadth and coverage can be found in Table 1 and Supp. Table 4. The sample “Almere 2018” was excluded from further analysis, as only 6% of the genome on one segment (L) had matching reads. All sequenced viral genomes are classified under the *Bandavirus zwieselense* species, sharing over 95% amino acid sequence identity in the RdRp with the type strain *Zwiesel* virus^71,72^. Only one genome of this species has been available, derived from a pooled sample of five positive bats found up to 200 km apart in Bavaria^52^. This consensus sequence likely reflects a mosaic of multiple viral genomes which can result in incorrect phylogenetic inferences. Moreover, within the S-Segment of this type strain at least 200 base pairs at the 5’ end of its NSs coding sequence are missing (Supp. Figure 8).

### Phylogenetic Analysis of B. zwiesel Genomes

To broadly contextualize our sequences within the *Bandavirus* genus, we constructed a maximum likelihood (ML) phylogenetic tree of the L segment nucleotide sequences using representative genomes from all known *Bandavirus* species, as well as the two unclassified *Malsoor virus* genomes.

All *Bandavirus zwieselense (B. zwiesel)* genomes cluster within the SFTSV-like Bandavirus clade with 100% bootstrap support (Figure 1). In this broader SFTSV-like clade, *Malsoor virus* and *B. zwiesel* together constitute a distinct lineage, identified thus far exclusively in bats. Moreover, the *B. zwiesel* sequences—encompassing the ancient (Penzlin 1919) genome—constitute a separate well-supported subclade (100% bootstrap support) distinct from *Malsoor virus*.

Focusing exclusively on our recovered genomes of the *Bandavirus zwieselense* species, we computed ML trees for each coding sequence (CDS) using MUSCLE 5.1^61^ and IQ-TREE^62^, as previously outlined. The phylogenetic analysis revealed two well-supported clades consistently found across all three segments. Furthermore, two sequences with minimal genetic divergence—collected one year and nine kilometers apart (Diekholzen and Bad Salzdetfurth)—stand out by forming a subclade with a long branch length in each segment. By contrast, two samples from the Netherlands (Groningen and Hoogezand), collected within the same year and located 16 km apart, do not feature a distinct subclade; in fact, except for the RdRp CDS tree, they appeared in different clades across all other CDS trees. Due to the limited number of samples and short coding regions, the support for deeper clades in the phylogeny is low, particularly for the NSs and N CDS on the S segment.

Panels b-e of Figure 2 present the ML phylogenetic trees for each CDS (RdRp, Gn/Gc, NSs, and N), with clades A and B highlighted. The samples in the map (Fig. 2a) carry a color code indicating their clade membership. All samples exhibit an identical clade membership regarding the two CDSs on the S segment (NSs:A, N:A or NSs:B, N:B). Across the ten samples, five distinct clade membership combinations across the segments were detected (Figure 2), suggesting multiple reassortment patterns.

**Fig. 2:**
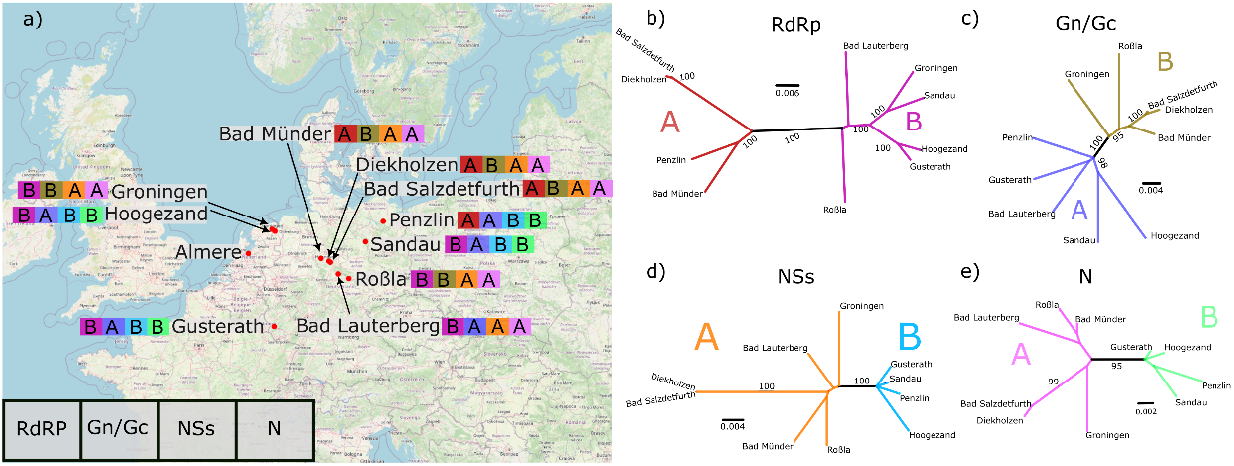
Locations of virus detection and phylogenetic relationships. **a)** Map indicating the location of positive tested bats as well as their color-coded composition of CDSs from the two major clades. Unrooted phylogenetic ML trees of the CDS of **b)** RdRp **c)** Gn/Gc **d)** NSs **e)** N. Each of the ML trees is color coded according to the two major clades A and B. The color coding highlights the composition of each viral genome in **a)**. The raw unmodified trees can be found in Supplementary Figures 4-7.

The samples from *Diekholzen, Bad Salzdetfuth* and *Bad Münder* share a common reassortment pattern (RdRp: clade A, Gn/Gc: clade B, NSs: clade A, N: clade A) and are geographically in close proximity. Despite being separated by several hundred kilometers, the samples from *Hoogezand, Sandau* and *Gusterath* also share a common reassortment pattern (RdRp: clade B, Gn/Gc: clade A, NSs: clade B, N: clade B). A third pattern (RdRp: clade B, Gn/Gc: clade B, NSs: clade A, N: clade A) is found for the samples from *Groningen* and *Roßla* that were collected hundreds of kilometers apart. Finally, the ancient sample *Penzlin* and the sample *Bad Lauterberg* possess a unique pattern of clade membership.

Given our heterochronous genomic data, we aimed to date the most recent common ancestor (MRCA) of all our sampled *B. zwiesel* viruses using the L segment. After confirming the presence of a clear temporal signal with TempEst v1.5.3^67^, we constructed a dated coalescent phylogeny using BEAST v1.10.4^68^, calibrating the strict molecular clock with tip dates. To additionally assess the impact of our ancient sample, we ran two analyses: once (***ancient+***) including the ancient sample and once (***ancient-***) excluding the ancient sample.

The estimates for the substitution rate and the year of the MRCA are shown in Figure 3. Both MRCA date distributions (Fig. 3a) are left-skewed, with the **ancient-** run exhibiting its mode in the year 1790, while the mode of the ***ancient+*** run is in the year 1470. Notably the 95% highest posterior density interval for the ***ancient-*** run extends until 1924, later than the 1919 collection date of the ancient sample from Penzlin. The substitution rate estimate (Fig. 3b) for the ***ancient+*** run is narrowly distributed at ~5 × 10^−5^ substitutions per site per year, in contrast with the ***ancient-*** run which has a broader distribution spanning nearly an order of magnitude.

**Fig. 3:**
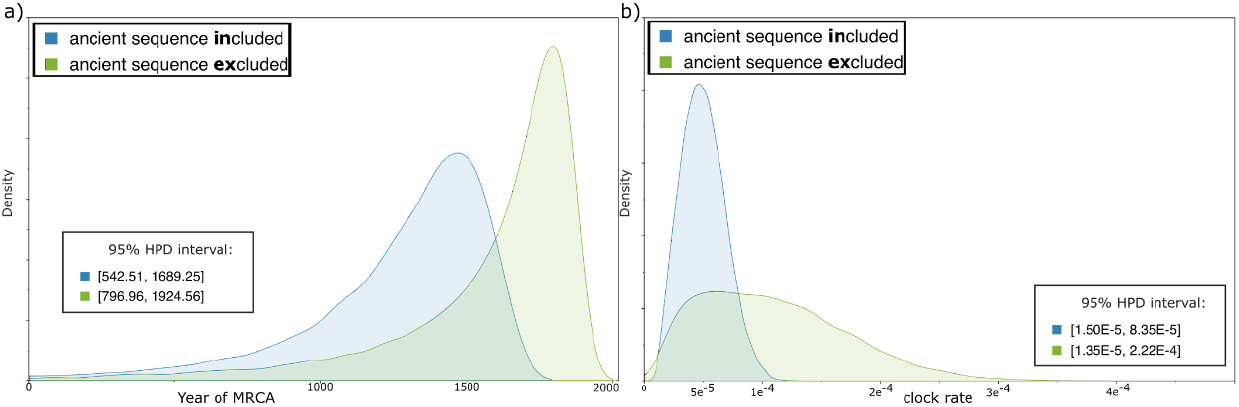
Marginal density plots of parameter estimates computed with BEAST v1.10.4^68^ using either the L segment of all our recovered *B. zwiesel* genome sequences or excluding the oldest sequence. a) Estimate of the year of the most recent common ancestor (MRCA). b) Estimate for the clock rate in substitutions/site/year.

### Functional characterization of the NSs protein

The immunomodulatory effects of the NSs protein of bandaviruses have been shown in multiple studies^20–24^. However, these properties of the NSs protein have not been investigated for *Malsoor virus* or *B. zwiesel*. To explore the functional properties of the *B. zwiesel* NSs protein, we examined its ability to suppress IFN induction and promote the formation of inclusion bodies.

To assess whether the *B. zwiesel* NSs protein can inhibit *Ifnb1* promoter activation, luciferase reporter assays were carried out in HEK293T cells. The reporter plasmids expressing luciferase under the control of the *Ifnb1* promoter were co-transfected with plasmids expressing MAVS, the key adaptor protein mediating IFN induction in response to RNA viruses. Additionally, varying amounts of plasmids expressing the NSs proteins from the Penzlin 1919 sequence (ancient), Bad Lauterberg 2010 sequence (recent), SFTSV or a negative control were co-transfected to compare their functionality. We were unable to also compare with the type strain Zwiesel virus, since its *NSs* gene is incomplete (Supp. Fig 8).

Both the ancient and recent NSs proteins demonstrated a strong, dose-dependent inhibition of MAVS-induced *Ifnb1* promoter activation (Figure 4 a). At the two highest amounts of plasmid, the inhibitory effect of both the ancient and recent untagged NSs is comparable to that of the human pathogenic SFTSV N-tagged NSs that was used as control. No relevant difference was observed between the inhibitory activity of the ancient and recent NSs protein variants.

**Fig. 4.**
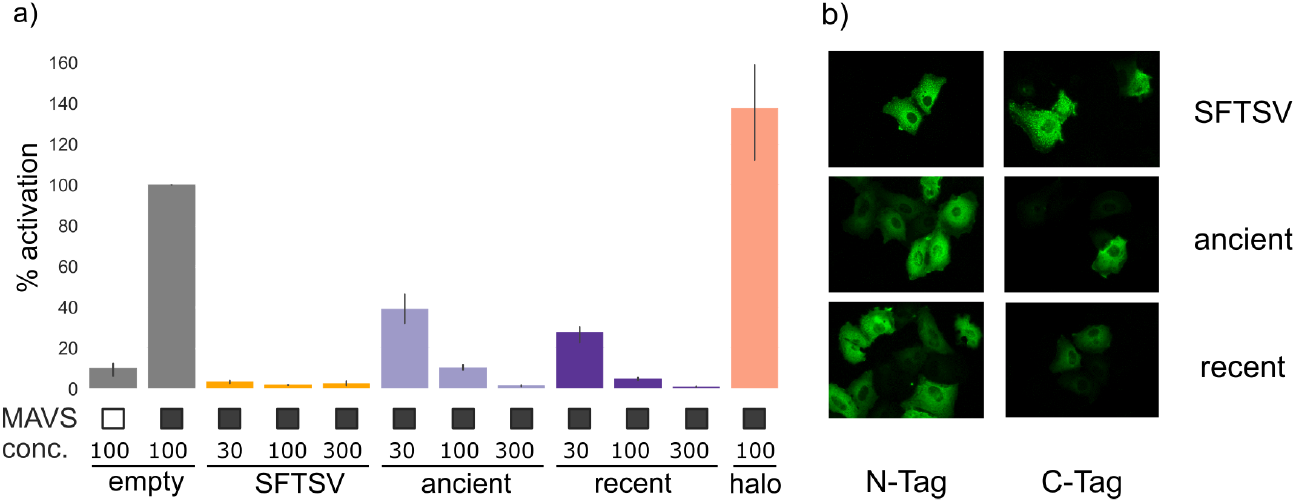
**a)** Firefly luciferase assay to measure *Ifnb1* promoter activation via MAVS overexpression in HEK 293T cells. Black empty or filled squares indicate whether MAVS was overexpressed or not. Below concentrations (conc.) of the respective plasmids are given in nanogram. Firefly luciferase activity was normalised to *Renilla* luciferase. The percentage of the activation of the empty vector is given for N-terminally 3×FLAG tagged SFTSV NSs, untagged Penzlin 1919 (ancient) NSs, untagged Bad Lauterberg 2010 NSs (recent) and N-terminally 3×FLAG tagged HaloTag as additional negative control. Error bars indicate estimated 95% confidence intervals. **b)** Huh7 cells were transfected with plasmids expressing either the N- or C-terminally 3×FLAG tagged Bad Lauterberg 2010 NSs (recent), Penzlin 1919 (ancient) NSs or SFTSV NSs and stained with mouse anti-FLAG antibody followed by goat anti-mouse IgG Alexa Fluor 488.

To evaluate whether the *B. zwiesel* NSs protein forms inclusion bodies similar to SFTSV NSs, we expressed N- and C-terminally 3×FLAG tagged *B. zwiesel* NSs in Huh7 cells for both the ancient and recent variants. The cellular distribution of the NSs proteins was assessed using fluorophore-conjugated anti-FLAG antibody staining. Cells transfected with plasmids expressing tagged SFTSV NSs protein were used as a positive control. As expected, the positive controls produced distinct inclusion bodies in the cells consistent with previous descriptions^20–22,27^ (Figure 4 b). In contrast, neither the ancient nor recent *B. zwiesel* NSs variants formed inclusion bodies. Instead, the NSs protein was evenly distributed throughout the cells, showing no evidence of localized concentration for neither the N-nor C-tagged *B. zwiesel* NSs proteins. Nevertheless, both tagged proteins retain their IFN induction inhibitory activity in HEK293T cells (Supp. Fig. 1 and 2).

## Discussion

The presented work highlights the potential of museum collections to contribute to RNA virus evolution studies. We report the successful extraction of viral RNA from an ethanol-preserved specimen after 100 years of storage. The recovered coding-complete viral genome enabled follow-up studies on virus presence in contemporary samples, phylogenetic analysis, and functional analysis of the encoded virus proteins. Our work extends on the findings of the first five *Bandavirus zwieselense* strains in Bavaria previously described by Kohl et al. ^52^, by reporting additional findings in midwest and northern Germany as well as the Netherlands. Furthermore, we identified two additional bat species (*P. pipistrellus, E. serotinus*) susceptible to infection and sequenced six coding-complete genomes as well as four partially complete genomes. Infection appears to be widespread across multiple organs as we detected viral RNA in liver, lung, feces, colon, brain, kidney and spleen. The related SFTSV is suspected to be primarily transmitted via ticks ^28,29^, making it conceivable that B. zwiesel is transmitted via the same route. A recent study focusing on the feeding patterns of bat-associated tick species concludes that bat-ticks may play a significant role as disease vectors, potentially facilitating pathogen transmission between bats and non-chiropteran hosts^73^.

The ten genome sequences we generated allowed the first phylogenetic analysis of the *Bandavirus zwieselense* species. Clade membership appears independent of sampling location and time (Figure 2) indicating substantial undersampling. As additional genomes are discovered, we expect more subclades to emerge that link clade membership with temporal and geographic patterns, as seen with samples from *Diekholzen* and *Bad Salzdetfurth*.

The clear division into two well-supported clades per segment led us to visualize putative reassortment patterns, hinting at frequent reassortment events within this species. Notably, these patterns are distributed across a wide geographic area, suggesting co-existing reassortants. The recovered genomes span 99 years, with the two oldest sequences collected in 1919 and 1999 and the most recent in 2018. Although the limited number of samples inevitably results in parameter estimates with relatively high uncertainty, the dated phylogeny constructed using the ancient sample (**ancient+**) provides a plausible estimate of the molecular clock rate. The ancient sample provides a lower bound on the dating of the MRCA. We see a clear violation of this lower bound for the dated phylogeny constructed without the oldest sample (***ancient-***). Even though the data for this phylogeny spanned 19 years, the derived molecular clock rate is too rapid, leading to an inaccurate 95% highest posterior density interval for the MRCA extending into the year 1924.

The estimated substitution rate under a strict clock model of ~5 × 10^−5^ substitution/site/year falls within the broad spectrum reported for RNA viruses (1 × 10^−2^ to 1 × 10^−5^)^74,75^. Considering that the most-likely mode of transmission is via ticks, arthropod viral substitution rates of ~1 × 10^−4^ that are reported for the related HRTV^76^ and the more distantly related *eastern equine encephalomyelitis virus (EEEV)*^*77*,*78*^ might be expected. Considering the time-dependent rate phenomenon^79^ which infers reduced evolutionary rates for longer observed time spans, the observed molecular rate seems plausible. Without virus isolates, any *de novo* assembly method carries the risk of generating chimeric sequences by incorrectly combining viral and non-viral fragments (e.g., from the host or associated microbes). The low sample quality did not allow for virus isolation and no readily available reverse genetics system exists. Hence, we employed the described overexpression approach to characterize one of the viral proteins and additionally validate the accuracy of our assembled genome. Our findings confirmed that the NSs proteins of both *B. zwiesel* strains are functional, exhibiting clear anti-IFN induction activity. A strong dose dependent inhibitory effect on IFN induction was observed, a feature that is known in SFTSV and HRTV, two human pathogenic bandaviruses^20,22,27,80^. This inhibitory effect was observed in a human kidney cell line, suggesting that the host immune components targeted by the *B. zwiesel* NSs are likely conserved across hosts and that its IFN suppression mechanism may extend to a broader range of species. Although the exact inhibitory process remains to be determined, the absence of inclusion bodies in fluorescently labeled *B. zwiesel* NSs-expressing cells (Figure 4 b), where N- and C-terminally tagged NSs proteins retained their inhibitory effect (Supp. Fig. 1 and 2), implies that the mechanism of IFN induction inhibition is distinct from that of SFTSV.

In future research, an unbiased sampling of contemporary specimens, coupled with serological surveys, could offer deeper insights into the distribution of *Bandavirus zwieselense* in Europe. Our demonstration of the feasibility of sequencing ancient viral genomes from museum samples may encourage further investigations, ultimately refining molecular clock rate estimates. Finally, the presented functional work raises the question of the precise mechanism that allows the NSs protein to inhibit IFN induction, a subject that should be investigated as well in bat cell lines to understand host-specific differences and assess the virus’s zoonotic potential.

## Data availability

The data for this study have been deposited in the European Nucleotide Archive (ENA) at EMBL-EBI under accession number PRJEB86147.

All code to reproduce the findings, figures and tables is available on github at the following URL: https://github.com/UdoGi/ancient_bunyavirus

## Acknowledgements

We gratefully acknowledge the Natural History Museum in Berlin, especially Frieder Mayer, Christiane Funk and Detlef Willborn for providing organs from ethanol-preserved bat specimens that were essential to this research, and Jim Robertson for providing the expression vector pI.18. Additionally, we thank our partners in the Netherlands: Peter Lina for species determination, and Lydia Dammers, Anja Sjoerdsma, Petra Vlaming, Antoinette van Wilgen and Stichting Vogelklas Karel Schot for providing bat carcasses.

