## SupplementaryMaterial for "A century-old museum sample reveals a bandavirus with modern day presence in northern European bats"

### Supplementary Materials

#### Supplementary Tables

| Federal state | # of specimens |
|---|---|
| Lower Saxony | 248 |
| Berlin | 204 |
| Saxony-Anhalt | 183 |
| Baden-Württemberg | 118 |
| unknown | 85 |
| Mecklenburg-Vorpommern | 59 |
| Rhineland-Palatinate | 26 |
| Brandenburg | 25 |
| Saarland | 21 |
| Hesse | 15 |
| Schleswig-Holstein | 7 |
| North Rhine-Westphalia | 5 |
| Thuringia | 4 |

**Supplementary Table 1:**
Origin of bat specimens by federal states in Germany

| Species | # of specimens | # tested postive |
|---|---|---|
| *Pipistrellus pipistrellus* | 355 | 6 |
| *Nyctalus noctula* | 177 | 0 |
| Not determined | 73 | 0 |
| *Eptesicus serotinus* | 51 | 1 |
| *Myotis nattereri* | 49 | 0 |
| *Myotis mystacinus* | 44 | 0 |
| *Myotis daubentonii* | 40 | 0 |
| *Plecotus auritus* | 39 | 0 |
| *Pipistrellus nathusii* | 30 | 0 |
| *Pipistrellus spec.* | 24 | 0 |
| *Vespertilio murinus* | 23 | 0 |
| *Nyctalus leisleri* | 18 | 0 |
| *Myotis myotis* | 17 | 0 |
| *Myotis brandtii* | 12 | 0 |
| *Plecotus austriacus* | 12 | 0 |
| *Pipistrellus pygmaeus* | 12 | 0 |
| *Myotis bechsteinii* | 6 | 0 |
| *Myotis myst/ bra /alca* | 5 | 0 |
| *Barbastella barbastellus* | 4 | 0 |
| *Myotis dasycneme* | 3 | 0 |
| *Eptesicus nilssonii* | 2 | 0 |
| *Myotis spec.* | 2 | 0 |
| *Plecotus spec.* | 1 | 0 |
| *Pipistrellus kuhlii* | 1 | 0 |

**Supplementary Table 2:**
Species composition of sampled contemporary bats in Germany

| Sampling round | Country | Family | Genus | Species | # of specimens | Min year | Max year |
|---|---|---|---|---|---|---|---|
| 1 | Cameroon | Pteropodidae | *Eidolon* | *helvum* | 6 | 1905 | 1937 |
| 1 | Cyprus | Pteropodidae | *Rousettus* | *aegyptiacus* | 3 | 1912 | 1912 |
| 1 | Germany | Vespertilionidae | *Pipistrellus* | *pipistrellus* | 8 | 1816 | 2000 |
| 1 | Indonesia | Molossidae | *Chaerephon* | *plicatus* | 2 | 1899 | 1926 |
| 1 | Israel | Pteropodidae | *Rousettus* | *aegyptiacus* | 5 | 1910 | 1910 |
| 1 | Italy | Vespertilionidae | *Nyctalus* | *noctula* | 1 | 1908 | 1908 |
| 1 | Madagascar | Pteropodidae | *Eidolon* | *helvum* | 1 | 1907 | 1907 |
| 1 | Peru | Phyllostomidae | *Desmodus* | *rotundus* | 1 | 1911 | 1911 |
| 1 | Poland | Vespertilionidae | *Nyctalus* | *noctula* | 2 | 1915 | 1919 |
| 1 | Rwanda | Pteropodidae | *Eidolon* | *helvum* | 3 | 1908 | 1908 |
| 1 | Switzerland | Vespertilionidae | *Nyctalus* | *noctula* | 1 | 1907 | 1907 |
| 2 | Germany | Vespertilionidae | *Eptesicus* | *nilssonii* | 9 | 1982 | 1986 |
| 2 | Germany | Vespertilionidae | *Pipistrellus* | *pipistrellus* | 28 | 1902 | 2000 |

**Supplementary Table 3:**
Composition of species and countries of origin of the samples from the Natural History Museum in Berlin. Two rounds of sampling took place. The first round covered various continents and species. The second targeted only Germany and the two bat species known to have been infected by *Bandavirus zwieselense*.

| Segment | Sample | Min coverage | Max coverage | Total mapped reads | Mean coverage | Breadth |
|---|---|---|---|---|---|---|
| L | Rossla 2011 | 286 | 6,226 | 96,238 | 2573.23 | 100.00 |
| M | Rossla 2011 | 24 | 8,004 | 118,702 | 4931.39 | 99.97 |
| S | Rossla 2011 | 37 | 8,004 | 36,603 | 3476.71 | 100.00 |
| L | Bad Salzdetfurth 2011 | 1 | 4,226 | 26,104 | 568.19 | 97.57 |
| M | Bad Salzdetfurth 2011 | 5 | 1,020 | 2,576 | 82.29 | 36.61 |
| S | Bad Salzdetfurth 2011 | 1 | 860 | 2,703 | 229.01 | 80.59 |
| L | Sandau 1999 | 1 | 2,517 | 13,095 | 245.43 | 90.76 |
| M | Sandau 1999 | 1 | 1,177 | 2,270 | 74.65 | 62.75 |
| S | Sandau 1999 | 3 | 1,572 | 4,599 | 308.89 | 88.26 |
| L | Gusterath 2010 | 4 | 4,665 | 77,135 | 1629.64 | 99.86 |

| | | | | | | |
|---|---|---|---|---|---|---|
| M | Gusterath 2010 | 2 | 8,000 | 78,955 | 3053.95 | 99.94 |
| S | Gusterath 2010 | 2 | 8,023 | 107,678 | 4902.19 | 100.00 |
| L | Bad Lauterberg 2010 | 50 | 8,044 | 2,041,742 | 7721.74 | 100.00 |
| M | Bad Lauterberg 2010 | 30 | 8,043 | 1,617,592 | 7540.12 | 100.00 |
| S | Bad Lauterberg 2010 | 160 | 8,057 | 665,167 | 7386.98 | 100.00 |
| L | Bad Münder 2012 | 300 | 8,032 | 681,046 | 7475.55 | 100.00 |
| M | Bad Münder 2012 | 39 | 8,072 | 1,592,158 | 7584.85 | 100.00 |
| S | Bad Münder 2012 | 88 | 8,055 | 459,195 | 7072.54 | 100.00 |
| L | Penzlin 1919 | 2 | 432 | 10,447 | 203.97 | 100.00 |
| M | Penzlin 1919 | 1 | 2,098 | 31,950 | 1162.82 | 100.00 |
| S | Penzlin 1919 | 1 | 701 | 5,896 | 423.06 | 100.00 |
| L | Hoogezand 2018 | 44 | 8,008 | 203,488 | 4800.49 | 100.00 |
| M | Hoogezand 2018 | 5 | 1,698 | 3,805 | 187.15 | 99.71 |
| S | Hoogezand 2018 | 1 | 8,018 | 34,508 | 2906.59 | 100.00 |
| L | Groningen 2018 | 1 | 424 | 5,362 | 131.77 | 99.95 |
| M | Groningen 2018 | 1 | 572 | 3,249 | 132.42 | 95.24 |
| S | Groningen 2018 | 2 | 395 | 1,270 | 105.45 | 95.24 |
| L | Diekholzen 2012 | 2 | 2,175 | 17,530 | 325.90 | 94.14 |
| M | Diekholzen 2012 | 1 | 1,613 | 5,855 | 205.92 | 84.72 |
| S | Diekholzen 2012 | 1 | 993 | 2,482 | 172.47 | 85.18 |

**Supplementary Table 4:**
Sequencing details for positive tested specimens. The sequencing reads across all organs of a specimen were merged for the purpose of genome assembly. Breadth: percentage of the genome covered by at least one read.

| | Almere 2018 | Hoogezand 2018 | Groningen 2018 | Gusterath 2010 | Diekholzen 2012 | Bad Salzdetfurth 2011 | Bad Münder 2012 | Roßla 2011 | Sandau 1999 | Bad Lauterberg 2010 |
|---|---|---|---|---|---|---|---|---|---|---|
| Liver | 1.7e0 | 1e2 | neg | 9.5e2 | neg | 1.1e2 | 1.7e3 | 1e1 | neg | 5.3e2 |
| Lung | neg | 3.7e2 | 2.3e2 | 7e2 | 2.7e0 | 2.1e2 | 8.5e2 | neg | neg | 1.1e3 |
| Feces | neg | 9.3e0 | neg | na | na | na | na | na | na | na |
| Nose wash | neg | neg | neg | na | na | na | na | na | na | na |
| Colon | neg | 3.7e2 | 4.9e1 | 2.4e1 | neg | neg | 1.1e1 | 7.7e1 | neg | 8.2e0 |
| Brain | 5.1e0 | 1.7e1 | neg | na | 6.7e1 | na | na | na | neg | na |
| Pharynx swab | neg | neg | neg | na | na | na | na | na | na | na |
| Rectal swab | neg | neg | neg | na | na | na | na | na | na | na |
| Kidney | neg | 3e1 | 9.1e1 | na | 1.8e2 | na | na | na | 4.5e2 | na |
| Spleen | neg | 1.1e3 | 2.4e1 | na | neg | na | na | na | neg | na |

**Supplementary Table 5:**
RT-qPCR details on positive tested specimens. The detected copies per microliter (cp/µl) are shown for each positive tested organ. neg: tested negative, na: sample not available.

| Country | Province | Species | Min year | Max year | # of specimens |
|---|---|---|---|---|---|
| Netherlands | North Brabant | *P. pipistrellus* | 2018 | 2020 | 36 |
| Netherlands | South-Holland | *P. pipistrellus* | 2018 | 2019 | 16 |
| Netherlands | Groningen | *P. pipistrellus* | 2018 | 2018 | 10 |
| Netherlands | Flevoland | *P. pipistrellus* | 2018 | 2018 | 7 |
| Netherlands | Utrecht | *P. pipistrellus* | 2018 | 2018 | 5 |
| Netherlands | North-Holland | *P. pipistrellus* | 2018 | 2018 | 4 |
| Netherlands | Drenthe | *P. pipistrellus* | 2018 | 2018 | 3 |
| Netherlands | Limburg | *P. pipistrellus* | 2018 | 2018 | 2 |
| Netherlands | Gelderland | *P. pipistrellus* | 2018 | 2018 | 2 |
| Netherlands | Friesland | *P. pipistrellus* | 2018 | 2018 | 1 |

**Supplementary Table 6:**
Summary of tested specimens from the Netherlands

### Supplementary Figures

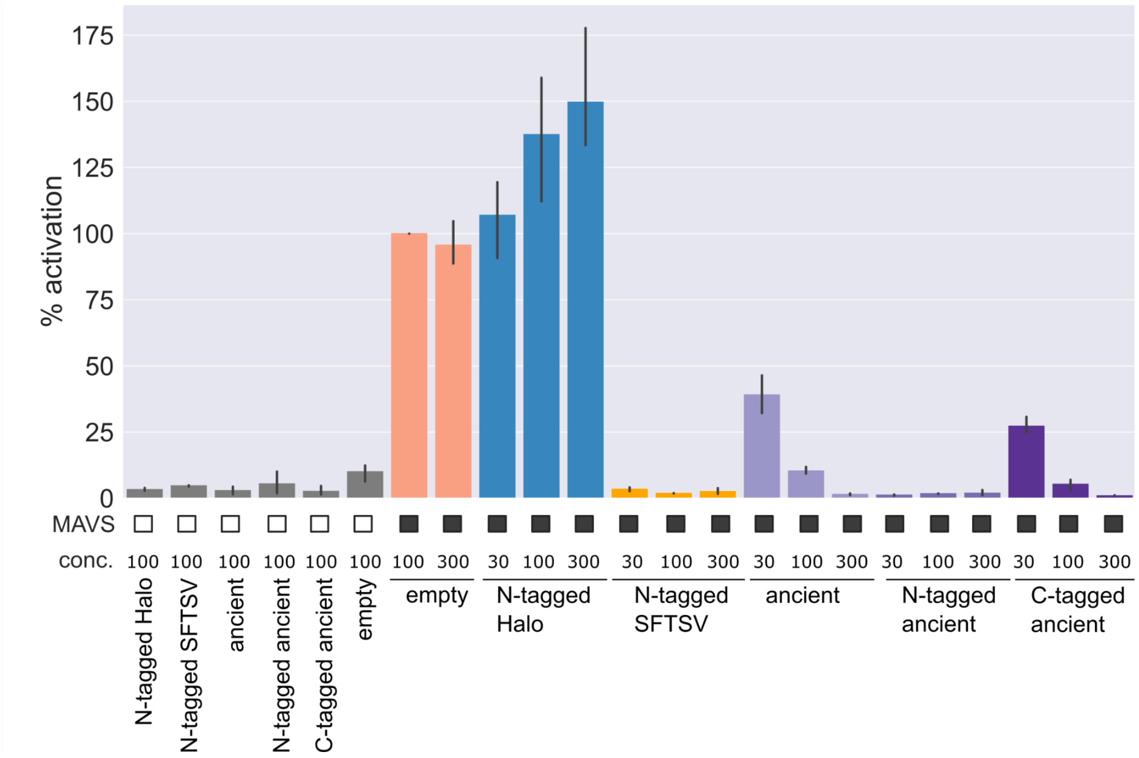


#### Supplementary Figure 1:

Firefly luciferase assay to measure *Ifnb1* promoter activation via MAVS overexpression in HEK 293T cells for the ancient Penzlin 1919 sequence. Black empty or filled squares indicate whether MAVS was overexpressed or not. Concentrations (conc.) of the respective plasmids are given in nanograms. Firefly luciferase activity was normalised to *Renilla* luciferase. The percentage of the activation of the empty vector (100 ng) is given for N-terminally 3×FLAG tagged HaloTag (N-tagged Halo) as additional negative control, N-terminally 3×FLAG tagged SFTSV NSs (N-tagged SFTSV),  untagged Penzlin 1919 NSs (ancient), N-terminally 3×FLAG tagged Penzlin 1919 NSs (N-tagged ancient), C-terminally 3×FLAG tagged Penzlin 1919 NSs (C-tagged ancient). Error bars indicate estimated 95% confidence intervals.

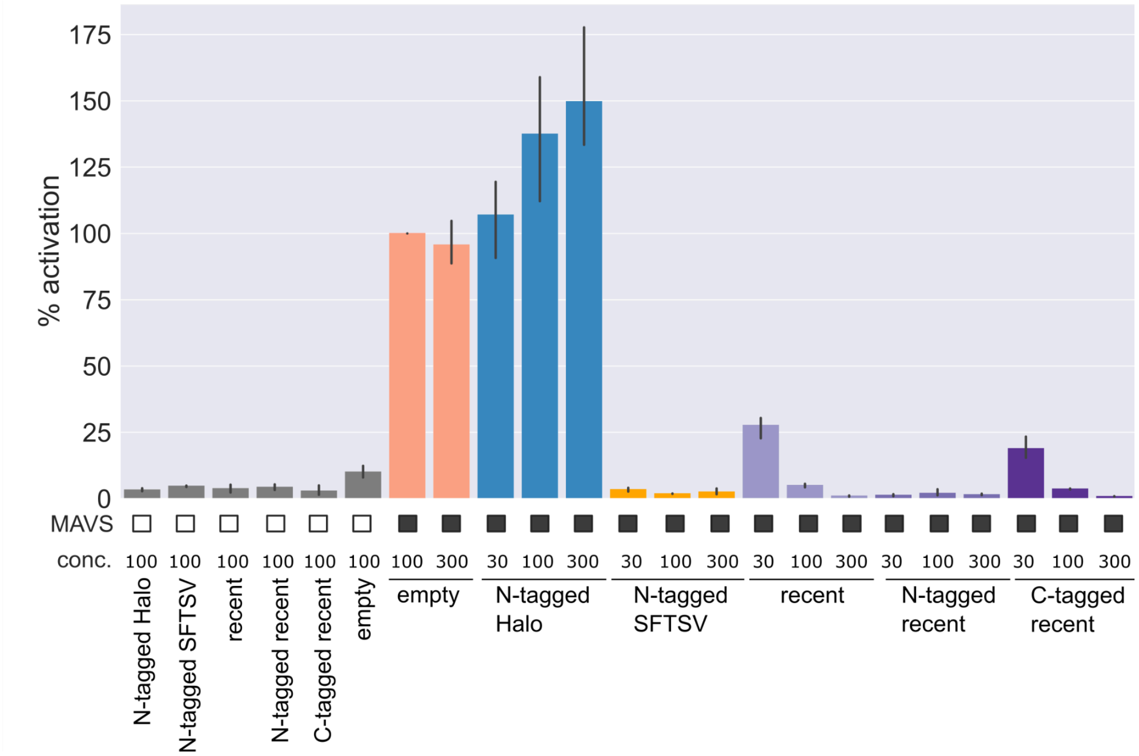


#### Supplementary Figure 2:

Firefly luciferase assay to measure *Ifnb1* promoter activation via MAVS overexpression in HEK 293T cells for the recent Bad Lauterberg 2010 sequences. Black empty or filled squares indicate whether MAVS was overexpressed or not. Concentrations (conc.) of the respective plasmids are given in nanograms. Firefly luciferase activity was normalised to *Renilla* luciferase. The percentage of the activation of the empty vector (100 ng) is given for N-terminally 3×FLAG tagged HaloTag (N-tagged Halo) as additional negative control, N-terminally 3×FLAG tagged SFTSV NSs (N-tagged SFTSV), untagged Bad Lauterberg 2010 NSs (recent), N-terminally 3×FLAG tagged Bad Lauterberg 2010 NSs (N-tagged recent), C-terminally 3×FLAG tagged Bad Lauterberg 2010 NSs (C-tagged recent). Error bars indicate estimated 95% confidence intervals.

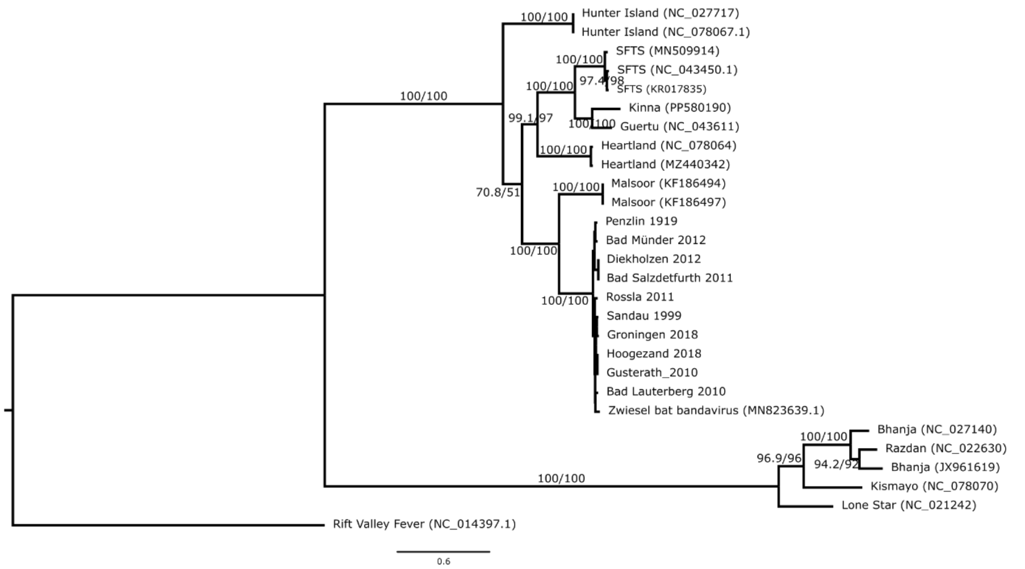


#### Supplementary Figure 3:

Phylogenetic relationships of L segment sequences of strains from the *Bandavirus* genus. The maximum likelihood tree was computed using IQ-TREE 2.3.6[62]. Rift Valley Fever Virus (NC_014397.1) was used as an outgroup for rooting. Bootstrap support values were computed using UFBoot[63] and SH-aLRT[64] and are both shown on the branches. Within the *B. zwiesel* clade no bootstrap support values are plotted.

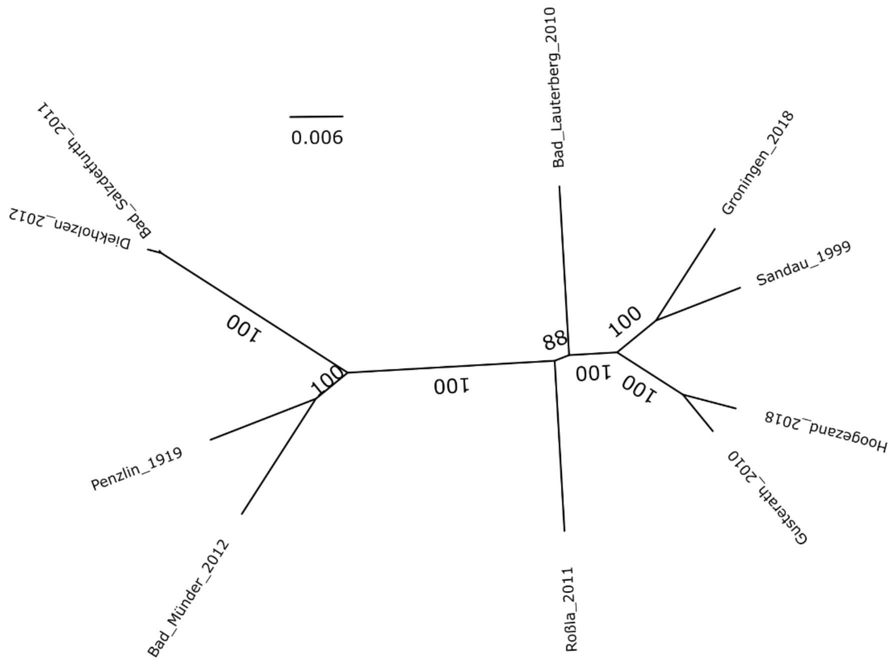


#### Supplementary Figure 4:

Unrooted Maximum Likelihood Tree of the RdRp CDS of our recovered genomes. Sequence alignment was performed with MUSCLE 5.1[61], followed by ML tree construction in IQ-TREE 2.3.6[62], which included a comprehensive model search and bootstrap value calculation with UFBoot[63] that are shown on the branches.

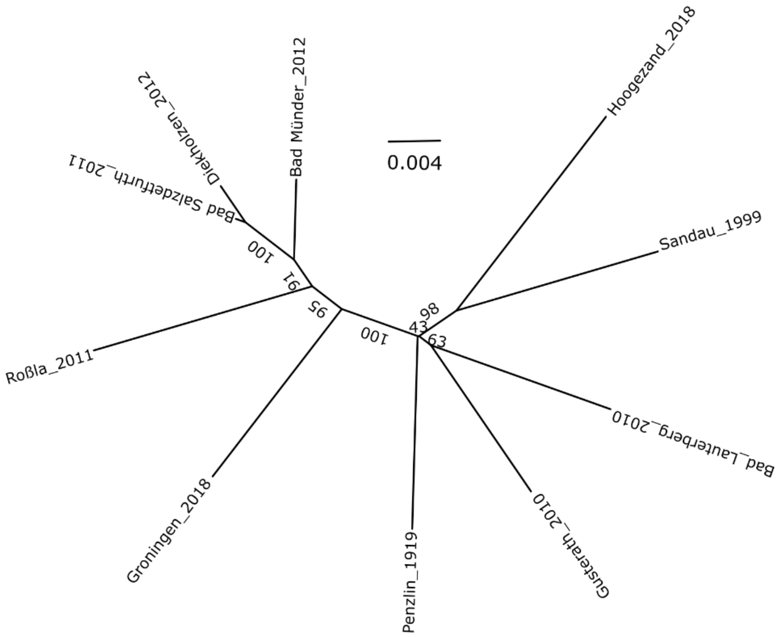


#### Supplementary Figure 5:

Unrooted Maximum Likelihood Tree of the Gn/Gc CDS of our recovered genomes. Sequence alignment was performed with MUSCLE 5.1[61], followed by ML tree construction in IQ-TREE 2.3.6[62], which included a comprehensive model search and bootstrap value calculation with UFBoot[63] that are shown on the branches.

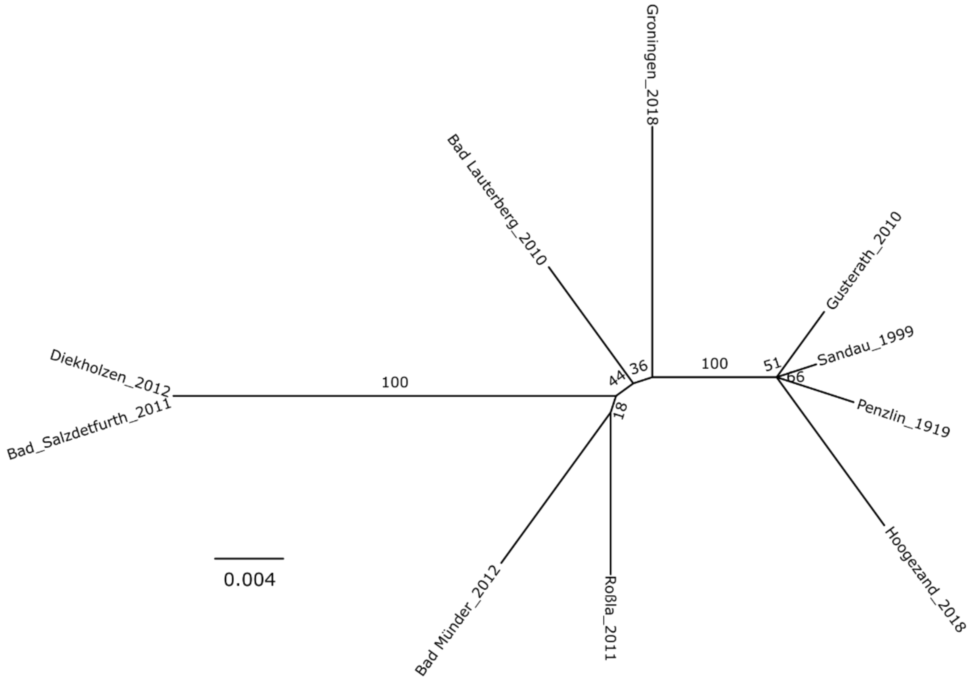


#### Supplementary Figure 6:

Unrooted Maximum Likelihood Tree of the NSs CDS of our recovered genomes. Sequence alignment was performed with MUSCLE 5.1[61], followed by ML tree construction in IQ-TREE 2.3.6[62], which included a comprehensive model search and bootstrap value calculation with UFBoot[63] that are shown on the branches.

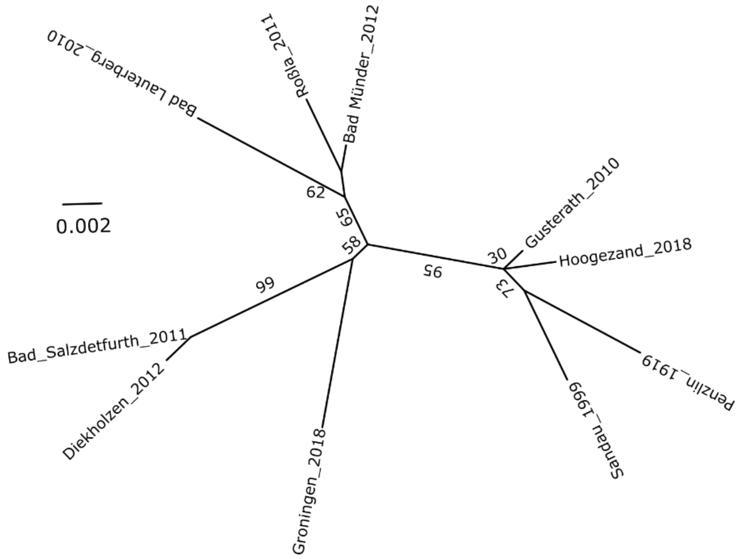


#### Supplementary Figure 7:

Unrooted Maximum Likelihood Tree of the N CDS of our recovered genomes.. Sequence alignment was performed with MUSCLE 5.1[61], followed by ML tree construction in IQ-TREE 2.3.6[62], which included a comprehensive model search and bootstrap value calculation with UFBoot[63] that are shown on the branches.

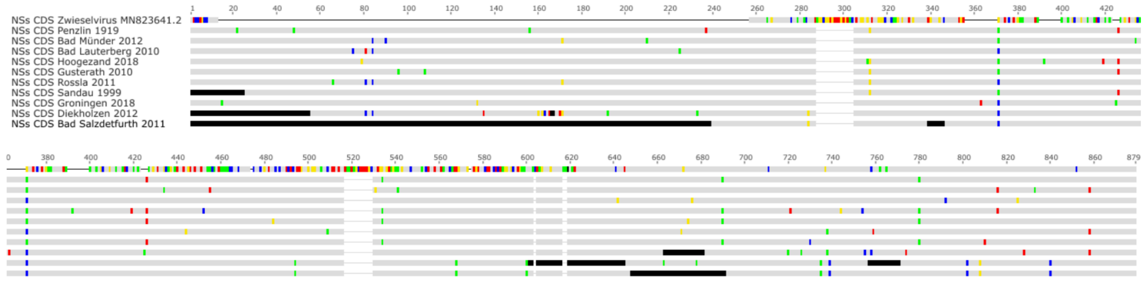


#### Supplementary Figure 8:

Sequence alignment (MUSCLE 5.1[61]) of the NSs CDS of our recovered genomes and Zwiesel virus (MN823641.2). The alignment is visualized in Geneious Prime® 2024.0.7, highlighting differences to the overall consensus sequence in color, black bars indicate missing sequence information. Zwiesel virus has a 606 bp NSs CDS, whereas our genomes span 846 bp, consistent with the 881 bp length observed in the related SFTS virus NSs CDS[81]. The first ~340 nucleotides of the Zwiesel virus sequence show no alignment to any of our 10 genomes, while the remaining ~260 nucleotides align with our recovered genomes.
